## Supplementary_Figs_Tables_S1andS2 for "Modulating gene expression and protein secretion in the bacterial predator *Bdellovibrio bacteriovorus*"

Mihajlovic *et al*.,

S1-S7 Figures page 2 - 8
S1-S2 Tables page 9
Supplementary References page 10

### Supplementary Figures

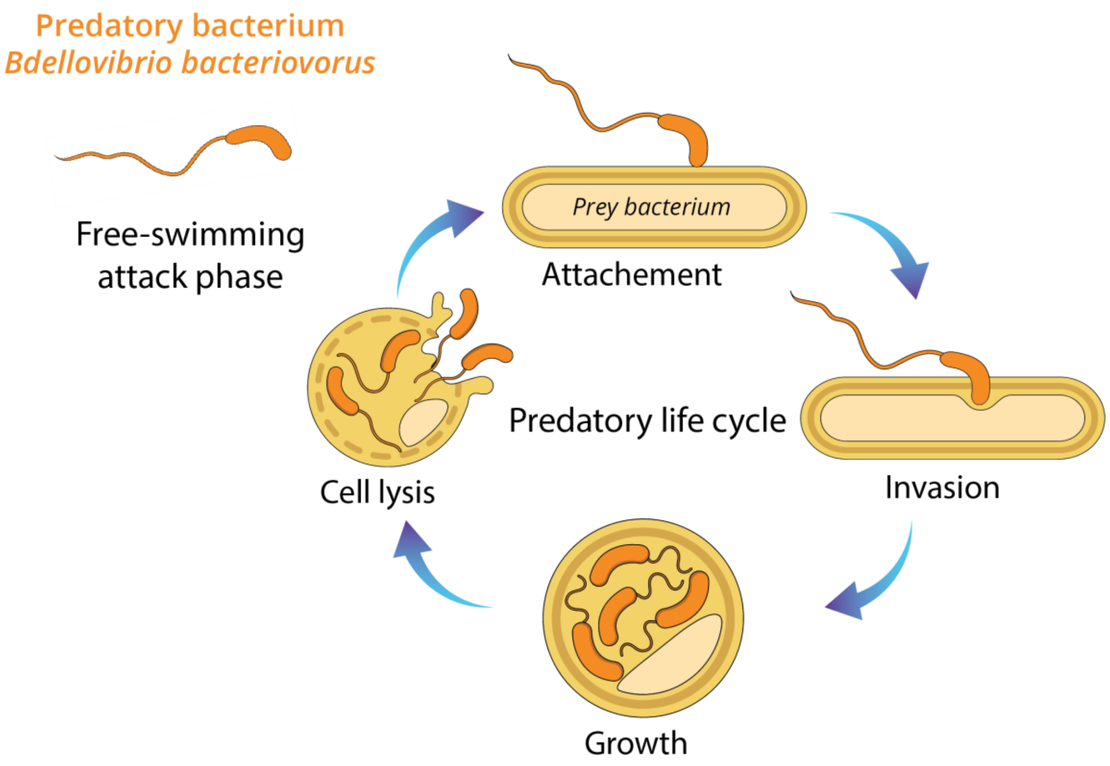


**S1 Figure. Overview of the *B. bacteriovorus* predatory life cycle.** During the free-swimming attack phase (AP), *B. bacteriovorus* swims towards and attaches to a Gram-negative bacterium (e.g. *Escherichia coli*). The predator then invades and multiplies within the prey periplasm, finally lysing the prey cell and releasing progeny back into the environment. These newly formed predator cells resume the attack phase, continuing their search for new prey bacteria.

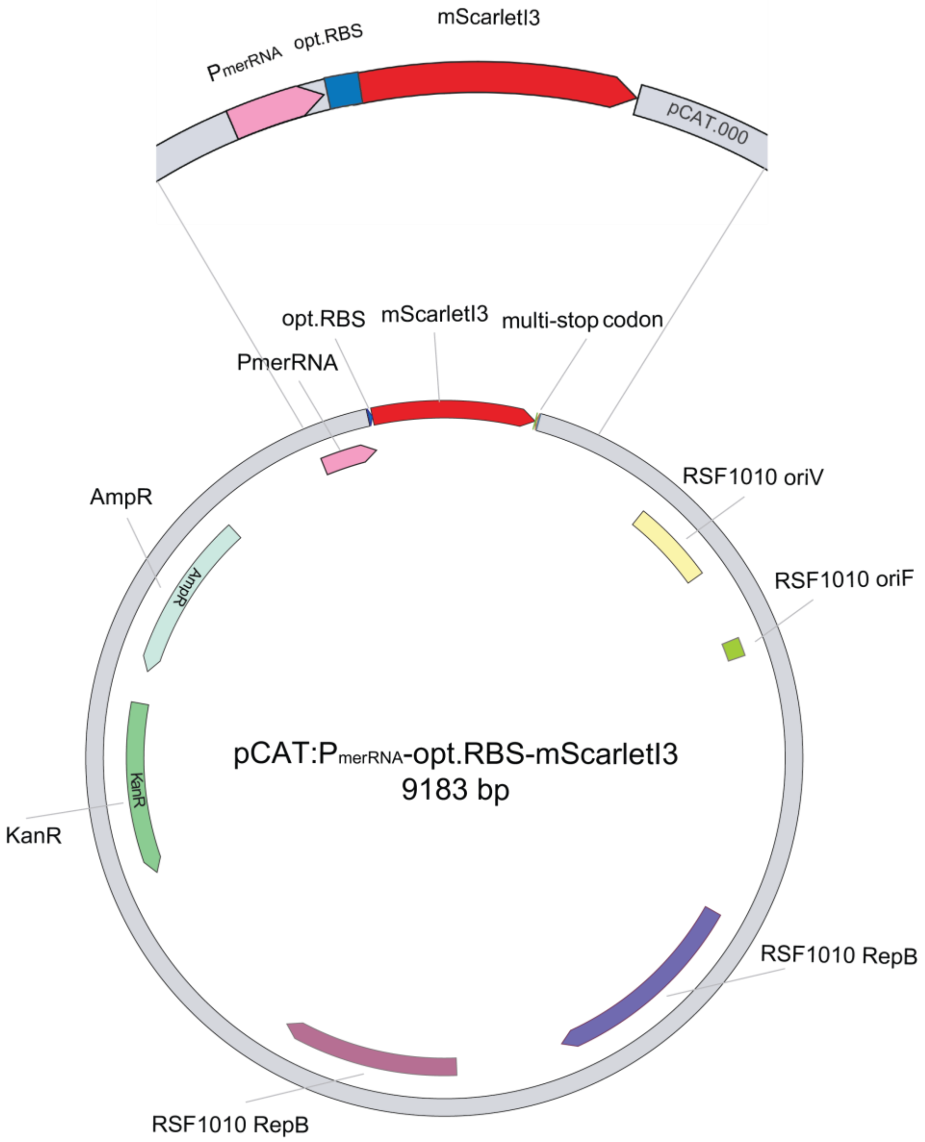


**S2 Figure.** Plasmid map of pLH-C1 (pCAT:P$_{merRNA}$-opt.RBS-mScarletI3), derived from pCAT.000 [1], showing the mScarletI3 reporter gene under the control of promoter P$_{merRNA}$ and an RBS sequence optimized for *B. bacteriovorus* (opt.RBS) [2]. Key features include kanamycin and ampicillin resistance cassettes (KanR, AmpR) for selection, as well as the RSF1010 origin of replication.

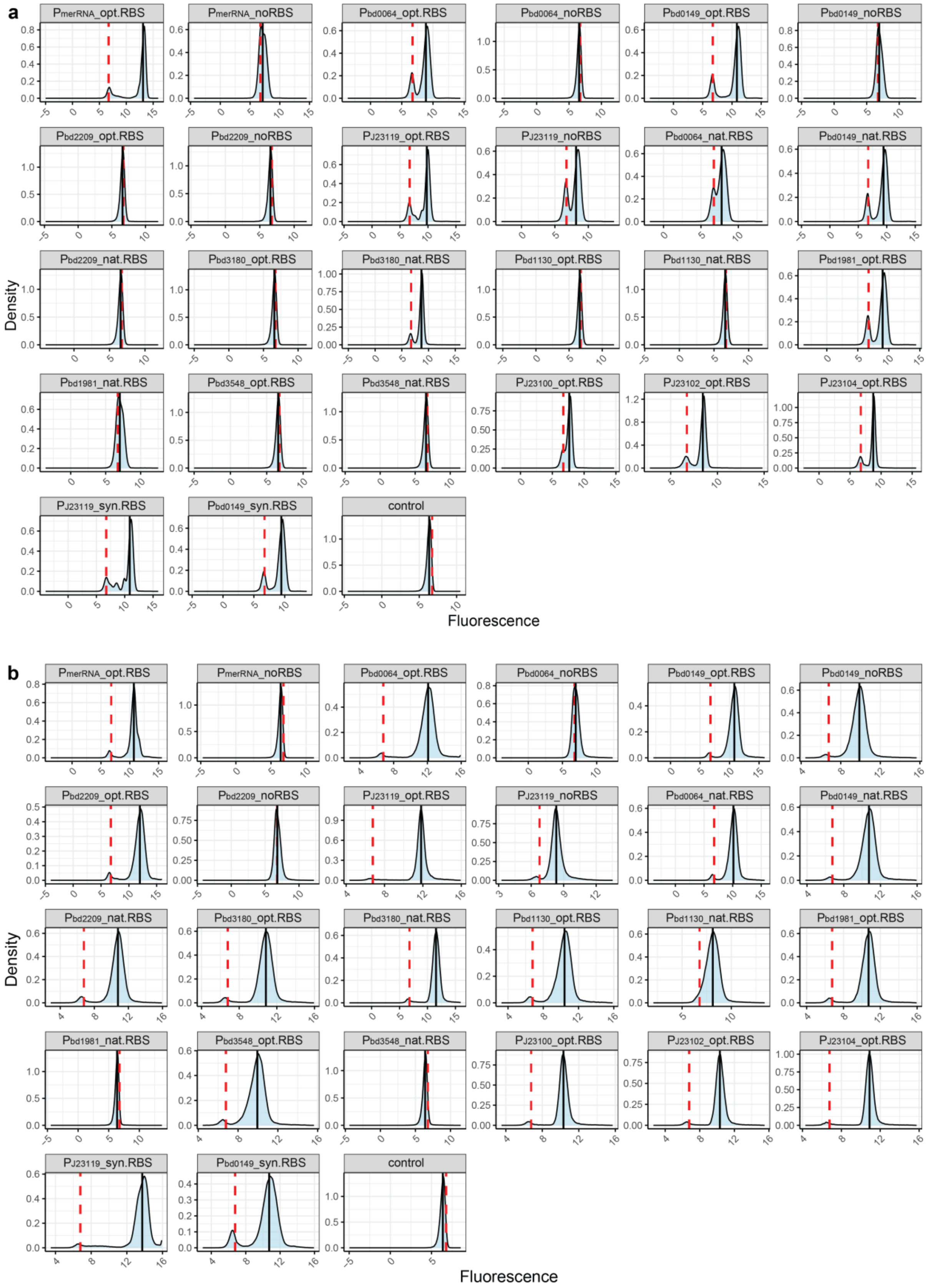


**S3 Figure. Overview of mScarlet fluorescence distributions in cells harbouring various pCAT.000-derived vectors in *B. bacteriovorus* AP (a) and *E. coli* S17-1 (b).** Thresholds (red, dashed line) indicate the 95$^{th}$ percentile of fluorescence intensity of the empty vector without reporter gene pCAT:P$_{merRNA}$-opt.RBS. Median fluorescence values are indicated by black lines. Panels show comparisons across different native and synthetic promoters combined with native (nat.RBS), optimized (opt.RBS), synthetic (syn.RBS), or no RBS (noRBS). Detailed plasmid description can be found in S5 Table.

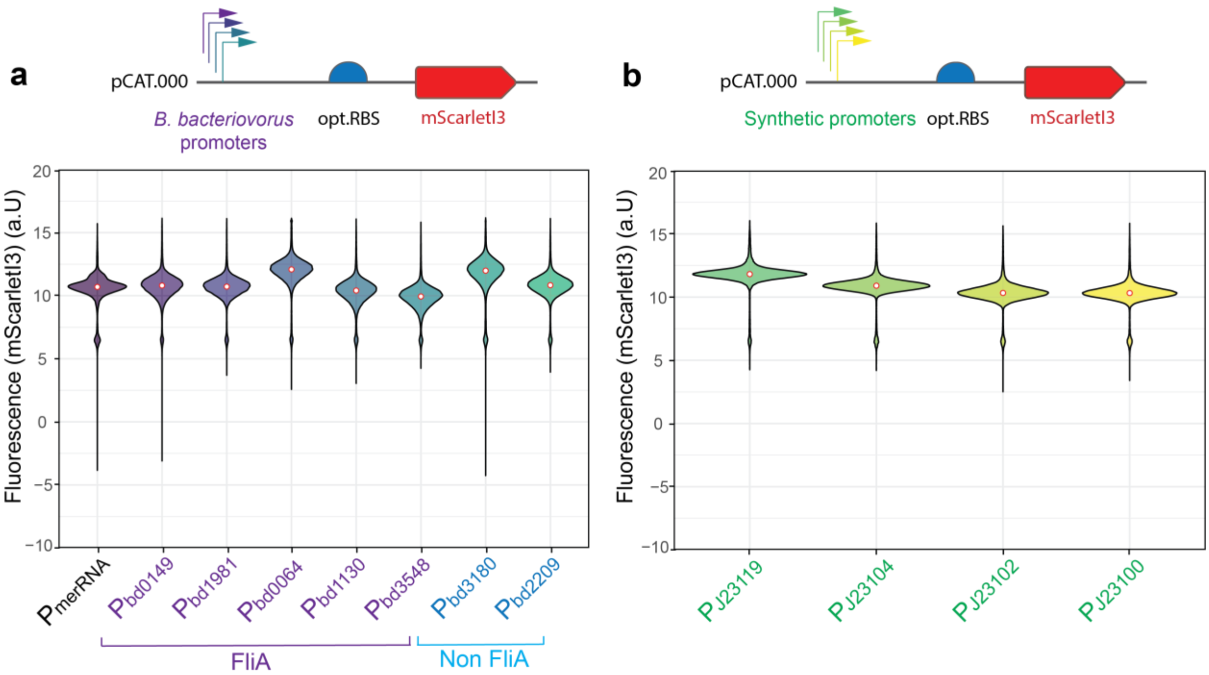


**S4 Figure. Native and synthetic promoters exhibit distinct mScarletI3 expression levels and variability in *E. coli* S17-1.** (**a**) Native *B. bacteriovorus* promoters, five with FliA motif ($P_{bd0149}$, $P_{bd1981}$, $P_{bd0064}$, $P_{bd1130}$, $P_{bd3548}$) and two without ($P_{bd3180}$, $P_{bd2209}$) were cloned upstream of mScarletI3 and introduced into *E. coli* S17-1. (**b**) Synthetic Anderson promoters ($P_{J23119}$, $P_{J23104}$, $P_{J23102}$, $P_{J23100}$) originally designed for *E. coli* drove varying levels of mScarletI3 expression under these same conditions. Each promoter was tested in a pCAT.000-derived plasmid with an optimized RBS upstream of mScarletI3. Fluorescence data was measured by flow cytometry. White dots in violin plots represent median fluorescence (density plots shown in S3b Fig). A second independent biological replicate yielded comparable results (see source data).

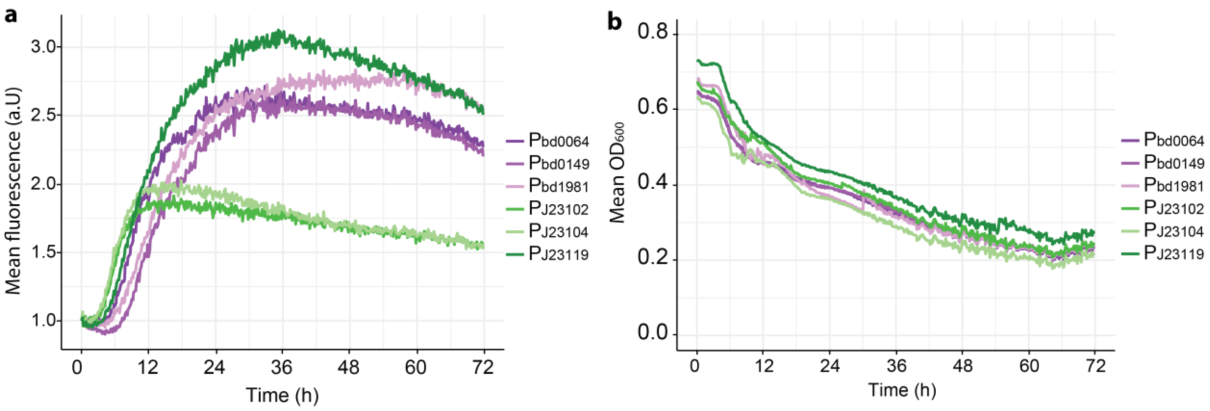


**S5 Figure. Extended temporal gene expression of selected synthetic and native promoters in *B. bacteriovorus* populations during predation over 72 hours.** (**a**) Overall gene expression level measured as mScarletI3 fluorescence at emission wavelength 517 nm during predation of *E. coli* S17-1 by *B. bacteriovorus* with native ($P_{bd0064}$, $P_{bd0149}$, $P_{bd1981}$) or synthetic ($P_{J23102}$, $P_{J23104}$, $P_{J23119}$) promoters expressed on pCAT.000-derived plasmids. For clarity of comparison, fluorescence starting values were adjusted to a common baseline across all samples. (**b**) Corresponding $OD_{600}$ changes over the extended predation period. Data represents the average of two biological replicates, each measured in two technical replicates.

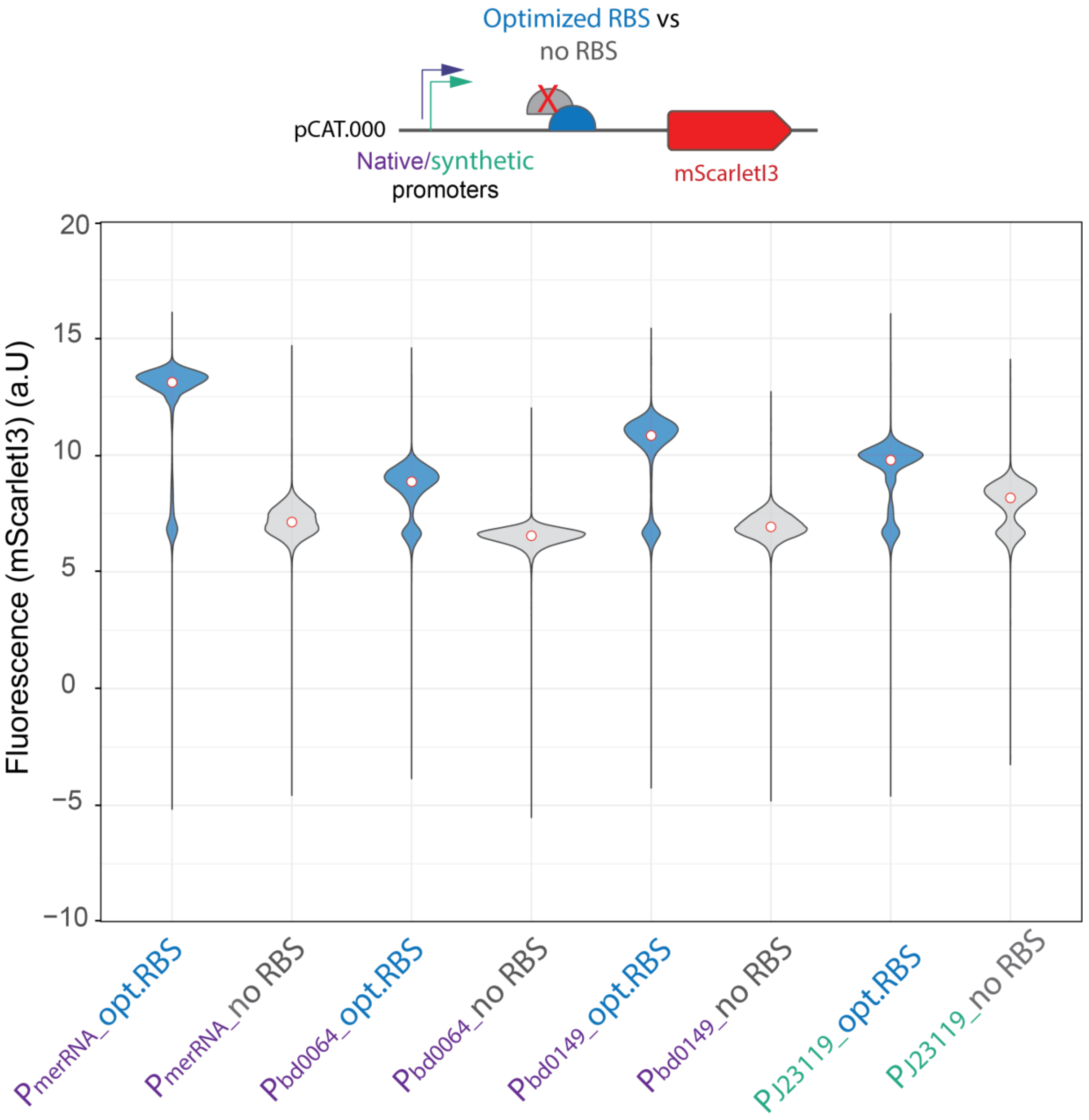


**S6 Figure. Removal of Ribosomal binding site (RBS) from *B. bacteriovorus* native promoters and synthetic promoters leads to low expression level of mScarletI3 in *B. bacteriovorus* AP populations.** Promoter strengths of *B. bacteriovorus* native promoters ($P_{merRNA}$, $P_{bd0064}$, $P_{bd0149}$) and the synthetic promoter $P_{J23119}$ were assessed at the population level in *B. bacteriovorus* AP using mScarlet-I3 fluorescence measured by flow cytometry. For all promoters, the *B. bacteriovorus* optimized ribosome binding site (opt. RBS, blue) was compared to the same construct lacking an RBS (no RBS, grey). White dots in violin plots represent median fluorescence (density plots shown in S3a Fig). A second biological and independent repeat of these measurements showed the same outcome (see source data).

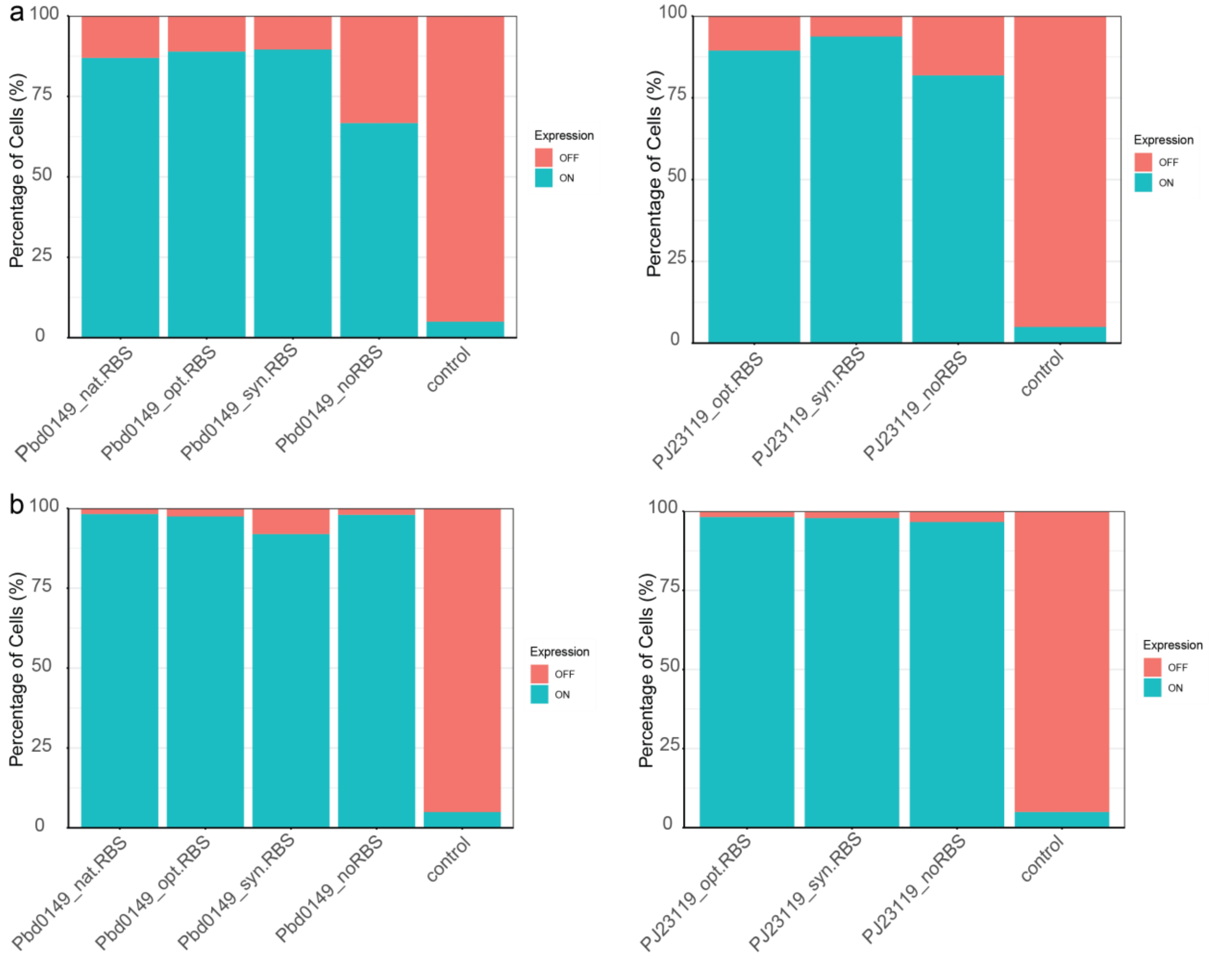


**S7 Figure. Effect of promoter and ribosome binding site (RBS) on gene expression levels in *B. bacteriovorus* AP (a) and *E. coli* S17-1 (b).** Bar plots represent the percentage of cells (%) exhibiting ON (cyan) or OFF (red) expression states, assessed using a fluorescent reporter gene of mScarletI3 on a pCAT.000-derived vector. Different combinations of the native promoter $P_{bd0149}$ (left) and the synthetic promoter $P_{J23119}$ (right) were tested with native (nat. RBS), optimized (opt. RBS), synthetic (syn. RBS), or no RBS (no RBS). Control represents cells harbouring empty vector without the reporter gene (pCAT:$P_{merRNA}$-opt.RBS). In general, the proportion of cells in ON state is larger in *E. coli* when RBS is absent compared to *B.bacteriovorus* AP (Figure S3), with exception of PmerRNA. Flow cytometry data was analysed as indicated in S7 Figure, where expression states were determined by gating cells based on fluorescence intensity thresholds set at the 95$^{th}$ percentile of the control. Most of this data is also shown as violin plots in Fig 4 andS4, S6 Figs.

**S1 Table. Overview of promoter regions used in this study.** Native promoter regions (violet stands for FliA regulated and light blue for non-FliA regulated, based on Karunker et *al*. [3] ), native sequence containing native RBS (grey underlined), optimized RBS (dark blue, underlined) with additional AAG as space holder (blue) to translational start site, synthetic RBS BBa_B0034 (orange). Abbreviations: not applicable (n/a).

| Native promoters | RBS | DNA Sequence 5`-> 3` | FliA regulation | Controlled gene |
|---|---|---|---|---|
| $P_{merRNA}$ | opt.RBS | AAAGAAACCTGTAACAGGTTTCTTTTTTCATCCAATTTCTAATTTTCAATTTCAAAAAGCCCCTCATCGAGGGGCTTTTTCTGTTTCAAGTCTGAACAAATATCCGTCTGTTTTCCCACGCTGATACGCAAATCTGACGTAATTTGAGAATCGCACGTGGTCCCCTTGAAAATAAAACTTTGGTCGAGCTATCCTGATTTTGTTGAAACCAGACTTAAGGAGGCAACAAGATG | no | merRNA |
| $P_{bd0149}$ | opt.RBS | GCGTCTCATTCTAATACAAGACCTAAAAACTTAAGAAACTCTGACTTTTGTCCGAAAAGGATATGGTAGGGGAAAGACTTAAGGAGGCAACAAGATG | yes | hypothetical protein |
| $P_{bd0149}$ | nat.RBS | GCGTCTCATTCTAATACAAGACCTAAAAACTTAAGAAACTCTGACTTTTGTCCGAAAAGGATATGGTAGGGGAAATTTTATG | yes | hypothetical protein |
| $P_{bd0149}$ | syn.RBS | GCGTCTCATTCTAATACAAGACCTAAAAACTTAAGAAACTCTGACTTTTGTCCGAAAAGGATATGGTAGGGGAAATCTAGAGAAAGAGGAGAAATACTAGATG | yes | hypothetical protein |
| $P_{bd1981}$ | opt.RBS | GGGACGGGGAAAATGTCTCAAAACTGGACCTGATGAAAAAACTCGTTTAAGCGTCCCTGCACCTCATTTATTCTATAAAGGTGGGGGAGCTGACTTAAGGAGGCAACAAGATG | yes | hypothetical protein |
| $P_{bd1981}$ | nat.RBS | GGGACGGGGAAAATGTCTCAAAACTGGACCTGATGAAAAAACTCGTTTAAGCGTCCCTGCACCTCATTTATTCTATAAAGGTGGGGGAGCTCATG | yes | hypothetical protein |
| $P_{bd0064}$ | opt.RBS | TTTTCAGAATTTTAATCCGAAAAGTTAAAAATCACCCGGACCCCCTATTAATTTTTGGTAAGTAATACCGAAAGGATAGTATCCAATTTAAGACTTAAGGAGGCAACAAGATG | yes | hypothetical protein |
| $P_{bd0064}$ | nat.RBS | TTTTCAGAATTTTAATCCGAAAAGTTAAAAATCACCCGGACCCCCTATTAATTTTTGGTAAGTAATACCGAAAGGATAGTATCCAATTTAAGCACAAGGAGTCTGACATG | yes | hypothetical protein |
| $P_{bd1130}$ | opt.RBS | CCTAAGAAAGAGACTCACATATTAAGGATTCTTCATCGTGAAAAACTCGTTTTTTCGGTCAAAAAACCGATAAGAAAGACATGAGAAACGGGACTTAAGGAGGCAACAAGATG | yes | putative YapH protein |
| $P_{bd1130}$ | nat.RBS | CCTAAGAAAGAGACTCACATATTAAGGATTCTTCATCGTGAAAAACTCGTTTTTTCGGTCAAAAAACCGATAAGAAAGACATGAGAAACGGAACGTCGCTTGTGTCATTCTTAGCACTCCTTTTTATG | yes | putative YapH protein |
| $P_{bd3548}$ | opt.RBS | CTTTTGGGATCTCGTGGAATATCACAACTTCCCGCCGGACAGCTCTTTACGATTCTTTGATGGAGTCCGAAGAGTCCACCAGACTTAAGGAGGCAACAAGATG | yes | hypothetical protein |
| $P_{bd3548}$ | nat.RBS | CTTTTGGGATCTCGTGGAATATCACAACTTCCCGCCGGACAGCTCTTTACGATTCTTTGATGGAGTCCGAAGAGTCCACCAATG | yes | hypothetical protein |
| $P_{bd3180}$ | opt.RBS | TACCGACACGCTGAAGTCGCTCCATTTCGTCTGGACCAAAGATGGTCCCGCCTTTTGTCCAAAAGGTGTAGAATGGGTGTGGCTCATAGGTGACTTAAGGAGGCAACAAGATG | no | hypothetical protein |
| $P_{bd3180}$ | nat.RBS | TACCGACACGCTGAAGTCGCTCCATTTCGTCTGGACCAAAGATGGTCCCGCCTTTTGTCCAAAAGGTGTAGAATGGGTGTGGCTCATAGGTGTTGAAATAGTGAGTGTTGTATAGAATGTGTTAAATCAAATTGAAAGATCAGGAGGATCACTATG | no | hypothetical protein |
| $P_{bd2209}$ | opt.RBS | CTGGCGGACCGGGTGGAGGAGTTAAGCCTTCGCCGCGACTCGATTCTGAAAAGAATGATTTATTTTAAAGTTAAGAAAGTTAATACAACGAGACTTAAGGAGGCAACAAGATG | no | hemolysin-type calcium binding protein |
| $P_{bd2209}$ | nat.RBS | CTGGCGGACCGGGTGGAGGAGTTAAGCCTTCGCCGCGACTCGATTCTGAAAAGAATGATTTATTTTAAAGTTAAGAAAGTTAATACAACGACGACGGCAAGTCTGTAGTTGGTGTTTGGGGGAAATTATG | no | hemolysin-type calcium binding protein |
| **Synthetic promoters** | | | | |
| $P_{J23119}$ | opt.RBS | TTGACAGCTAGCTCAGTCCTAGGTATAATGCTAGCGACTTAAGGAGGCAACAAGATG | no | n/a |
| $P_{J23119}$ | syn.RBS | TTGACAGCTAGCTCAGTCCTAGGTATAATGCTAGCGACTTCTAGAGAAAGAGGAGAAATACTAGAAGATG | no | n/a |
| $P_{J23104}$ | opt.RBS | TTGACAGCTAGCTCAGTCCTAGGTATTGTGCTAGCGACTTAAGGAGGCAACAAGATG | no | n/a |
| $P_{J23102}$ | opt.RBS | TTGACAGCTAGCTCAGTCCTAGGTACTGTGCTAGCGACTTAAGGAGGCAACAAGATG | no | n/a |
| $P_{J23100}$ | opt.RBS | TTGACGGCTAGCTCAGTCCTAGGTACAGTGCTAGCGACTTAAGGAGGCAACAAGATG | no | n/a |

**S2 Table. Overview of the Amino acid and DNA sequences used forSec-dependent signal peptides fused to NanoLuc reporter gene used in this study.** Cleavage sites were predicted with SignalP 6.0 [4] and are indicated with a red slash.

| N-terminal sec-dependent signal | Amino acid sequence | DNA Sequence 5`-> 3` |
|---|---|---|
| ss_Bd2269$_{1-22}$ | MKFNVFALIVSVLFATSAQA/ER | ATGAAATTCAACGTGTTTGCACTCATCGTATCAGTGCTTTTCGCGACATCAGCTCAGGCAGAGCGC |
| ss_Bd0468$_{1-23}$ | MKRPISLALSALTLTASLAHA/QE | ATGAAACGCCCTATTTCTTTGGCACTGTCCGCGCTGACTTTGACTGCTTCCCTGGCTCACGCCCAGGAG |
| ss_Bd0120$_{1-22}$ | MKKTTLIAALILLSSAAAHA/GD | ATGAAAAAAACCACACTGATTGCAGCCTTGATCCTGCTAAGCTCTGCAGCTGCACACGCCGGTGAC |
| ss_Bd2692$_{1-20}$ | MKRALLVGAVLFGSQAFA/GE | ATGAAACGTGCATTACTAGTTGGTGCGGTTCTGTTTGGTTCTCAGGCGTTCGCTGGCGAA |
